# Similar destabilization of neural dynamics under different general anesthetics

**DOI:** 10.1101/2025.08.21.671540

**Authors:** Adam J. Eisen, Alexandra G. Bardon, Jesus J. Ballesteros, André M. Bastos, Jacob A. Donoghue, Meredith K. Mahnke, Scott L. Brincat, Jefferson Roy, Yumiko Ishizawa, Emery Brown, Ila R. Fiete, Earl K. Miller

**Affiliations:** The Picower Institute for Learning and Memory, Massachusetts Institute of Technology, Cambridge, MA 02139, USA; McGovern Institute for Brain Research, Massachusetts Institute of Technology, Cambridge, MA 02139, USA; Department of Brain and Cognitive Sciences, Massachusetts Institute of Technology, Cambridge, MA 02139, USA; Department of Psychology, Ruhr-Universität-Bochum, 44801 Bochum, Germany; Department of Psychology, Vanderbilt University, Nashville, TN 37235, USA; Vanderbilt Brain Institute, Vanderbilt University, Nashville, TN 37235, USA; Beacon Biosignals, Boston, MA 02114, USA; Department of Anesthesia, Critical Care, and Pain Medicine, Massachusetts General Hospital, Harvard Medical School, Boston, MA 02139, USA; Division of Sleep Medicine, Harvard Medical School, Boston, MA 02115, USA

## Abstract

Different classes of anesthetics can induce unconsciousness despite acting through distinct biological mechanisms. This raises the possibility that they produce a convergent effect on the dynamics or temporal evolution of neural population activity. To explore this, we analyzed intracortical electrophysiological recordings during infusions of propofol, ketamine, and dexmedetomidine, using a rigorous method to estimate dynamical stability. We found that all three anesthetics, despite their molecular differences, similarly affect cortical states by destabilizing their dynamics. This destabilization matched the slower recovery from sensory perturbations and longer stimulus-induced autocorrelation times observed during the anesthetic infusions. The destabilization was also reflected predominantly in lower-frequency ranges, linking it to the well-known increase in low-frequency power during anesthesia. Finally, destabilization closely tracked real-time fluctuations in consciousness. Together, these findings suggest that cortical destabilization may be a shared neural correlate of anesthetic-induced unconsciousness, offering a mechanistic explanation for low-frequency oscillations observed during anesthesia.

## Introduction

A remarkable property of anesthetic-induced unconsciousness is that it is achieved by different anesthetics through distinct pathways of molecular influence. Anesthesia thus presents an exciting lens through which to explore consciousness,^1–5^ as seemingly disparate low-level mechanisms generate related impacts at the level of arousal and awareness.^6^

Propofol boosts inhibition through GABA_A_ (γ-aminobutyric acid type A) receptors, significantly altering cortical dynamics.^7–12^ Ketamine has been characterized as primarily a disinhibition of inhibitory interneuron projections onto cortex.^3,13–16^ Ketamine is an N-methyl-D-aspartate (NMDA) antagonist, and has been shown to specifically target NMDA receptors on inhibitory interneurons.^3,13–15^ The resulting decrease in excitatory input to these inhibitory interneurons thus disinhibits the interneurons’ projections. Dexmedetomidine, in contrast, is an agonist of α_2_-adrenergic receptors (primarily in locus coeruleus).^17^ Dexmedetomidine is thought to alter arousal through a pathway involved in sleep, in which agonizing α_2_-adrenergic receptors results in reduced firing of noradrenergic neurons and subsequently reduced inhibitory output from locus coeruleus.^18–20^ Despite the variety of molecular actions among these three anesthetic agents, all result in loss of consciousness, and all three evoke stereotyped changes in the frequency spectrum of brain activity, most notably boosting low frequencies.^6,7,12,19,21–29^ It is unclear how these dissimilar molecular mechanisms can all lead to similar changes in both arousal and neural oscillations.

In this work, we take a dynamical systems perspective towards understanding anesthetic-induced unconsciousness, characterizing the effects of different anesthetics on ongoing activity in cortex. We apply a novel and rigorous method (delayed linear analysis for stability estimation, or DeLASE) to quantify the *dynamical stability* of cortical activity over time, before and during anaesthetic administration and during recovery.^30^ D*ynamic stability* (henceforth stability) is a fundamental concept in dynamical systems theory and control. It is a measure of the robustness of a dynamical system, characterizing the rate at which the system recovers from disturbances (e.g., inputs, distractions, random fluctuations in activity) and returns to its baseline state (Figure 1A).

**Figure 1:**
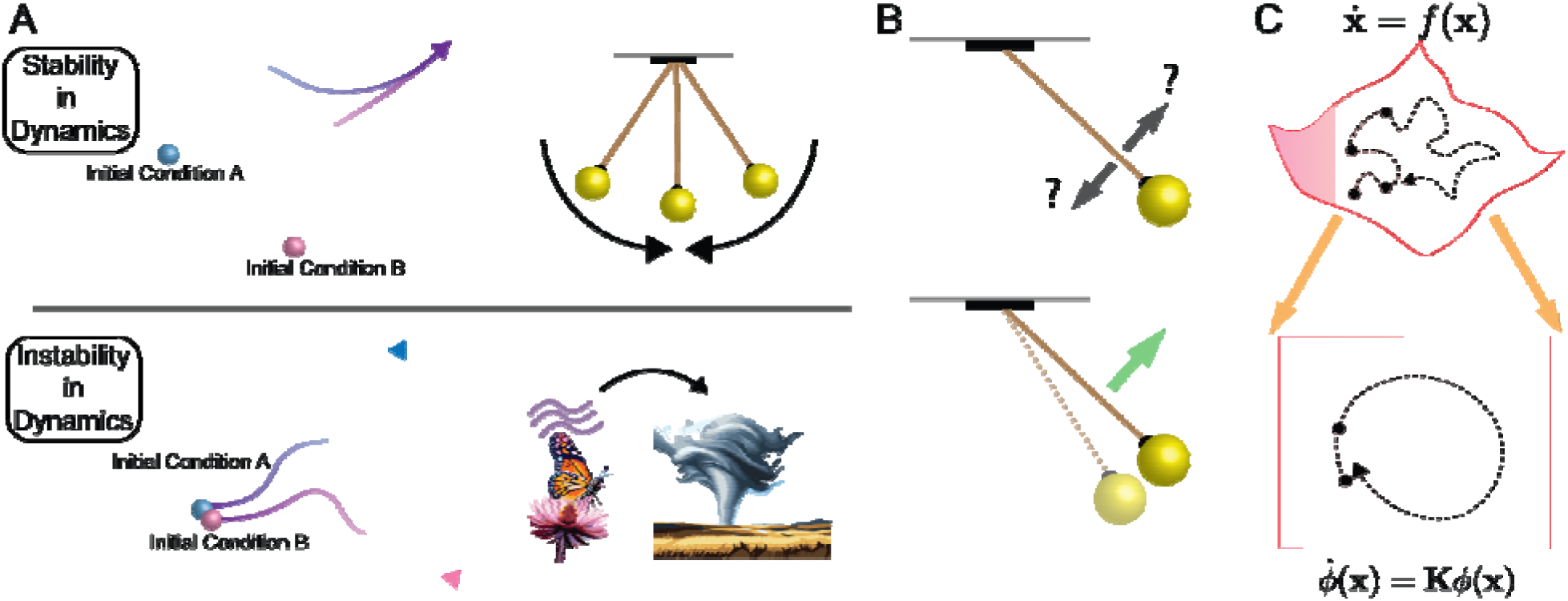
A dynamical systems approach to understanding neural dynamics during anesthesia. (A) Stability and instability in dynamics. (Top) Stable systems (e.g., a damped pendulum) show trajectories converging from different initial conditions. (Bottom) Unstable systems (e.g., weather) exhibit diverging trajectories even from similar initial conditions, illustrating sensitivity to small perturbations. (B) Trajectory history for prediction. Knowing only current position (left) is insufficient for prediction; trajectory history (right) reveals the influence of unobserved variables (e.g., velocity) on observed states. (C) Koopman operators enable linear representations of nonlinear dynamics. A nonlinear system’s dynamics on a manifold (top) can be linearly represented in a higher-dimensional space using techniques like time-delay embedding (bottom).

Through this approach, we seek global signatures of loss of consciousness that arise in the brain independent of local molecular actions. This approach - characterizing activity changes that are signatures of the high-level transition from consciousness to its absence - sheds light on the necessary mechanisms of consciousness that are disrupted by anesthetic agents. It was demonstrated previously through the application of DeLASE that propofol destabilizes cortical neural dynamics.^30^

DeLASE leverages the fact that a nonlinear dynamical system can be recovered from partial observations by considering a signal’s history simultaneously at multiple delay lags.^31–54^ For instance, if we observe a snapshot of a pendulum’s position but not its velocity (a partial observation), we cannot predict its next state (Figure 1B). By including the position history, the upcoming dynamics of the pendulum become clear. DeLASE uses a decorrelated, low-rank representation of the signal’s history to estimate a dynamical systems model of the data using linear regression.^55–57^ The field of Koopman operator theory provides guarantees that these models can represent nonlinear dynamics, despite their linearity (Figure 1C).^58–72^ Finally, to construct the stability estimates, DeLASE uses an algorithm that discounts the weight of earlier time delay lags to estimate the solutions of a differential equation representation of the model.^73,74^ These solutions correspond to the (inverse) characteristic timescale at which the system will respond to a perturbation. Thus, large timescale values indicate the system will decay quickly in response to perturbation. Smaller values indicate it will have a more extended—and thus more unstable—response.

Using intracortical electrophysiology from non-human primates (NHPs), we found that propofol, ketamine, and dexmedetomidine all induce a destabilization of cortical neural dynamics. This destabilization was evident in how the brain responds to sensory perturbations during the unconscious state, and was reflected across the low-frequency spectrum. We suggest that the stereotypical increase in low-frequency power during anesthetic-induced unconsciousness reflects the brain’s slower recovery times in the more unstable state.

## Results

### All three anesthetic agents destabilize neural dynamics

Local field potential (LFP) recordings from the cortex revealed that the anesthetics had destabilizing effects on cortical activity. We applied DeLASE^30^ to LFPs recorded from four NHPs before, during and after infusion of each anesthetic in different sessions. There were two NHPs used for each anesthetic. For the propofol sessions, electrodes were placed in the ventrolateral prefrontal cortex, frontal eye fields, posterior parietal cortex, and auditory cortex. For the ketamine and dexmedetomidine sessions, electrodes were placed bilaterally in the ventrolateral and dorsolateral prefrontal cortex. Instability values were computed in 15 second windows tiled across each session. DeLASE measures stability by computing how long it takes a system to recover from perturbation. We analyzed the instability values from the upper 10% of the distribution of characteristic roots (Figure 2). We used the upper 10% because they have the most impact on dynamic stability.

**Figure 2:**
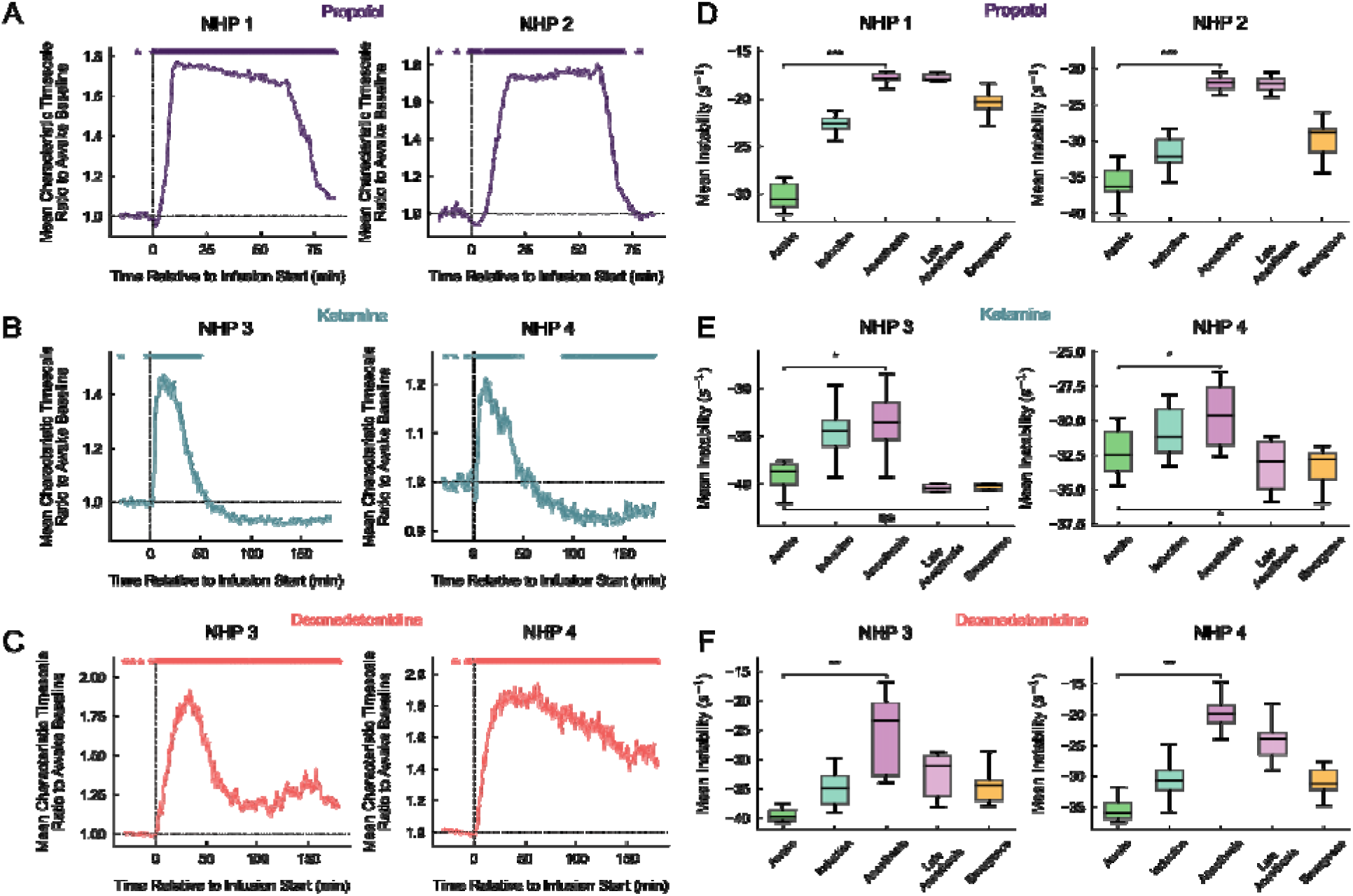
Propofol, ketamine, and dexmedetomidine destabilize neural dynamics. Data are represented as mean ± SEM, with results averaged over sessions for each NHP. (A, B, C) The mean timescales of response normalized to the awake baseline for propofol (A), ketamine (B) and dexmedetomidine (C). The left column corresponds to one NHP involved in the data collection and the right column corresponds to the other. The awake baseline was computed for each session by taking the geometric mean of the timescales associated with the characteristic roots across all windows in the awake section of the session. Then, for each window, the geometric mean of the ratio of the timescales to the awake baseline was computed. Horizontal dotted line corresponds to a ratio of 1 - i.e., when the mean timescale is equivalent to the awake baseline. The vertical dotted line corresponds to the start of the anesthetic infusion. Stars at the top of each plot indicate significance above 1, with n=the number of sessions recorded for each NHP. In each case, neural activity was robustly destabilized. (D, E, F) The mean instability grouped by epoch of the session. The Awake epoch was the part of the session before the infusion began. The Induction epoch was defined as 15 minutes following the start of infusion. The Anesthesia epoch was defined as the period from 15 minutes post infusion to 45 minutes post infusion.

Figure 2 shows instability normalized to the awake baseline state (Figure 2A-C) as well as raw instability values (Figure 2D-F). The normalized values show a clear destabilization that began shortly after infusion of anesthetic doses of each agent. The propofol sessions, which involved a continuous infusion, showed a sustained destabilization over the entire 60 minute infusion. The Anesthesia epoch was destabilized compared to the Awake epoch (p < 0.001 for both NHPs, one-sided Wilcoxon signed-rank test). The ketamine and dexmedetomidine sessions involved an initial bolus infusion. They predictably showed a rapid initial destabilization followed by a gradual decline, matching the metabolization of a bolus infusion. Again, for both ketamine and dexmedetomidine, the Anesthesia epoch was destabilized compared to the Awake epoch (p < 0.05 for all combinations of agent and NHP, one-sided Wilcoxon signed-rank test). Ketamine was unique in that it appeared to produce greater stability later in the session after initially destabilizing cortex (p < 0.05 for NHP3 and NHP4 considered together, one-sided Wilcoxon signed-rank test).

### Recoveries to sensory perturbation matched changes in dynamic stability

In the previous section, we established that all three agents destabilized (and in ketamine, ultimately stabilized) cortical neural dynamics. We now investigate how this destabilization changed recovery from perturbation using auditory sensory stimulation.

More stable systems recover faster from perturbation than less stable systems. An example of this is two systems in which different masses hang from identical springs (Figure 3A). With a smaller mass the system is more stable. Thus, when perturbed with identical force, the spring with the smaller mass initially moves more but then decays back to its resting state more quickly than the larger mass (Figure 3B). There are longer autocorrelation times, as perturbations linger for longer (Figure 3B).

**Figure 3:**
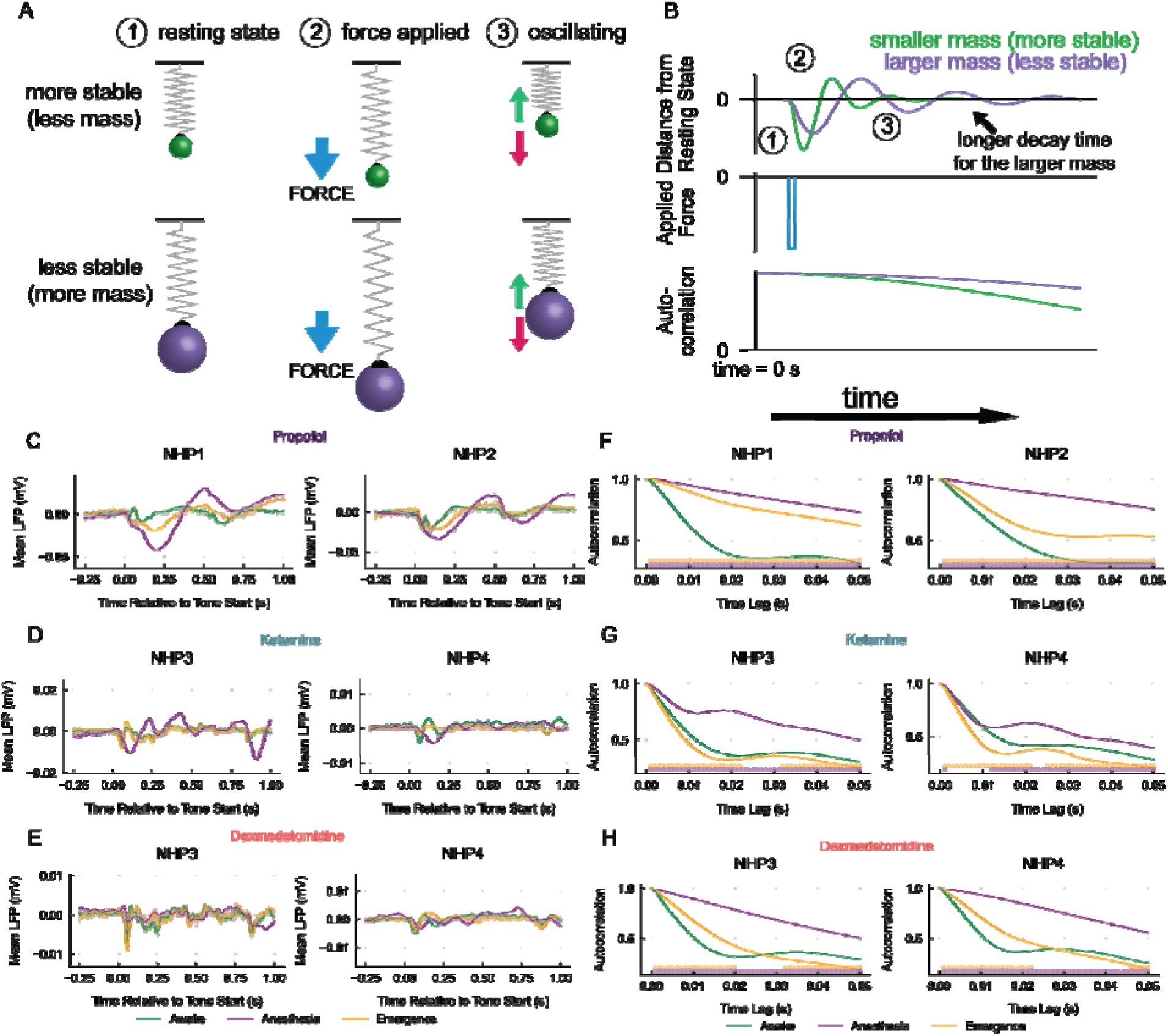
Destabilized dynamics explain sensory responses to tones during anesthetic unconsciousness. Data are represented as mean ± SEM, with results averaged over sessions for each NHP. (A) Mass-spring oscillators with spheres of different masses hanging from identical springs. (1) Spheres perturbed upward by equal force (2) oscillate back to the resting state (3). Smaller mass (green) decays faster than larger mass (purple). (B) Plots of the relevant time series for the example in (A). (C, D, E) The LFP response to auditory stimuli for propofol (C), ketamine (D), and dexmedetomidine (E). Response was averaged across electrodes and trials within a session, then the mean was computed across sessions. Translucent curves correspond to response averages from single sessions. Green curve is the Awake epoch, purple is the Anesthesia epoch, and orange is the Emergence epoch. Auditory stimuli were 500 ms for propofol (C), and auditory oddball stimuli consisting of five 150 ms spaced by 50ms for ketamine and dexmedetomidine (D and E, respectively). (F, G, H) Mean response autocorrelation functions for propofol (F), ketamine (G), and dexmedetomidine (H). Autocorrelation functions were computed separately for each electrode and averaged across electrodes and trials from all sessions. Green curve is the Awake epoch, purple is the Anesthesia epoch, and orange is the Emergence epoch. Stars on the bottom indicate whether Anesthesia is significantly different than Awake (purple stars) and whether Emergence is significantly different than Awake (orange stars). See also Figure S1.

We analyzed the neural responses to sensory perturbation during Awake, Anesthesia, and Emergence epochs. In propofol sessions, NHPs were periodically exposed to (500 ms) auditory tones throughout the session. In ketamine and dexmedetomidine sessions, NHPs underwent an auditory oddball task (five 50 ms tones spaced by 150 ms). We averaged neural responses across trials and electrodes for each session to obtain a time-series representation of the recovery from sensory to perturbation. We then averaged across sessions to obtain a single time-series.

The recoveries from sensory perturbation matched predictions from destabilized systems. For all the agents, the sensory responses showed slower recovery from perturbation during the Anesthesia epoch compared to Awake (Figure 3C-E, Figure S1). During the Emergence epoch, there was a difference between agents. For both propofol and dexmedetomidine, perturbation recovery remained slower compared to the Awake epoch, but faster than the Anesthesia epoch (Figure 3C,E). By contrast, for ketamine, after an initial period of slower recovery from perturbation, there was a prolonged period of shorter recovery (i.e., increased stability) relative to Awake (Figure 3D).

These results were quantified using an autocorrelation function of response to the tones, computed over 1 second of data following the tone onset, and averaged across trials and electrodes within each session. For all three agents, the Anesthesia epoch was characterized by longer autocorrelations values (Figure 3F-H, significance computed using the two-sided Wilcoxon signed-rank test). During emergence from propofol and dexmedetomidine, the autocorrelation values dropped relative to Anesthesia but were still higher compared to Awake epochs. By contrast, during emergence from ketamine, autocorrelation values were smaller than those in the Awake epoch.

Higher vs lower autocorrelation values are directly related to higher vs lower frequency LFP dynamics. In the next section, we investigate the relationship between stability and spectral power.

### Dynamic stability is reflected across lower frequencies

Longer autocorrelation values during destabilization implies an increase in lower frequency power. To analyze this relationship, for each session we computed the (Spearman) correlation between the time series of mean instability (DeLASE) values and the time series of average power in each frequency band. The frequency bands we considered were delta (0.1 - 4 Hz), theta (4 - 8 Hz), alpha (8 - 12 Hz), beta (12 - 30 Hz), and gamma (30 - 100 Hz).

Destabilization following all three agents correlated positively with delta, theta, and alpha power (Figure 4A-C, p < 0.01 for all combinations of NHP, agent, and band, two-sided Wilcoxon signed-rank test). Once again, there was a contrast between propofol and dexmedetomidine vs ketamine. For propofol and dexmedetomidine, gamma power was negatively correlated with destabilization (p < 0.01 for all combinations of NHP and agent, two-sided Wilcoxon signed-rank test). For ketamine it was positively correlated with destabilization (p < 0.05 for both NHPs, two-sided Wilcoxon signed-rank test), reflecting increases in gamma power during ketamine-induced unconsciousness^23^. Results were weaker and not consistent for beta (Figure 4A-C).

**Figure 4:**
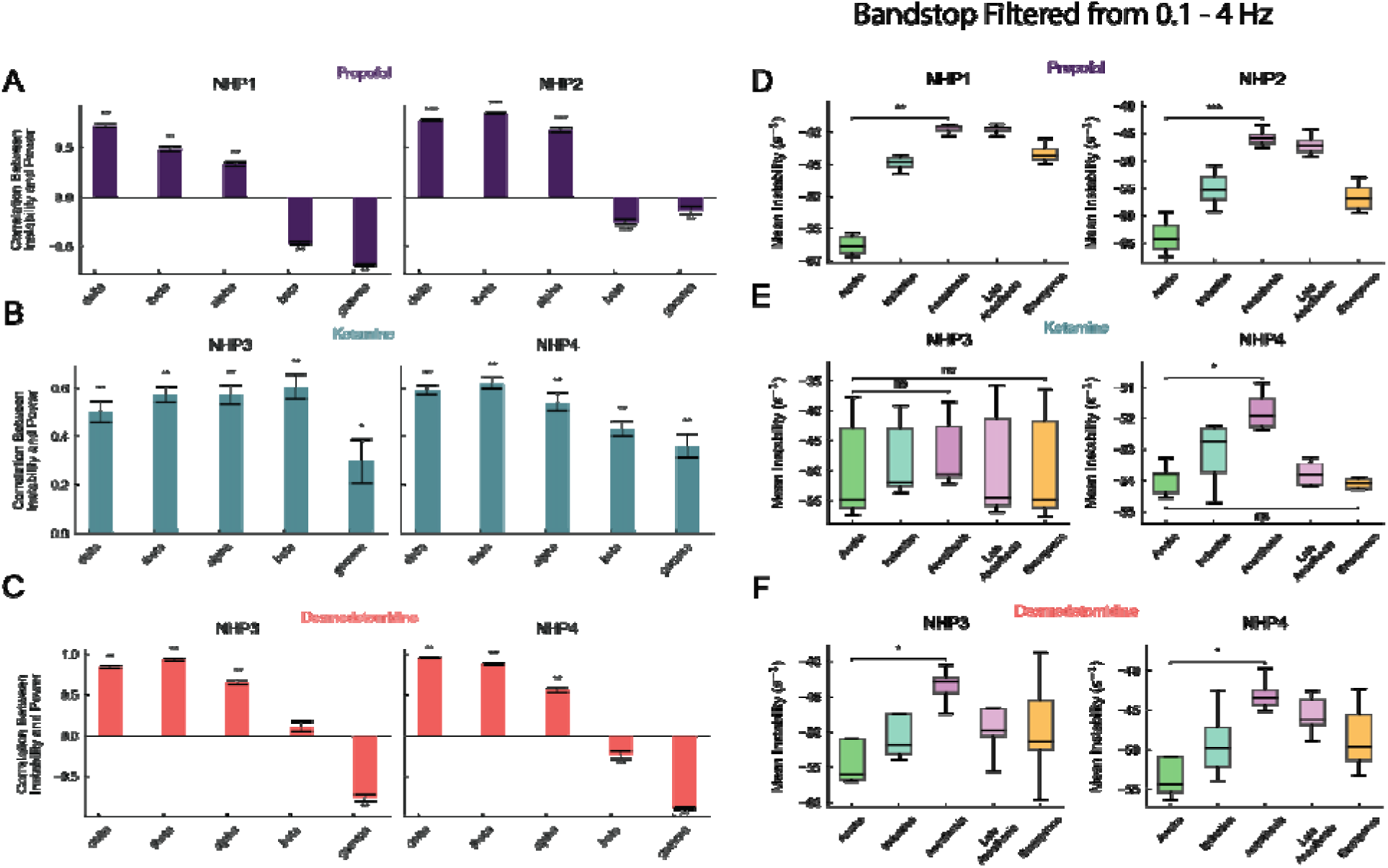
Changes in dynamic stability during anesthetic unconsciousness are reflected across the frequency spectrum. Data are represented as mean ± SEM, with results averaged over sessions for each NHP. (A, B, C) Mean correlation between instability values and band power for propofol (A), ketamine (B), and dexmedetomidine (C). Correlation values were averaged across sessions. Stars indicate significant differences from 0. (D, E, F) Mean instability values computed from data after removing the delta (0.1 - 4 Hz) frequency range for propofol (A), ketamine (B), and dexmedetomidine (C). See also Figure S2.

A shift to delta in the cortex is a hallmark of loss of consciousness due to anesthesia (Figure S2).^6,7,12,21–24^ To determine if destabilization was primarily driven by delta power, we applied a band-stop filter to all LFPs, effectively removing the 0.1 - 4 Hz frequency range from these signals. We then reapplied DeLASE to every session, and found that the destabilization results were similar (Figure 4D-F). This affirmed that information about dynamic stability is contained in a wide range of the frequency spectrum, not just in the delta frequencies.

### Instability tracks degrees of wakefulness in real time

In the dexmedetomidine sessions, the NHPs were fluctuating between responsiveness and unresponsiveness as they gradually emerged from the anesthetic infusion. Dexmedetomidine is in fact known to produce an arousable sedation state, in which subjects can move easily between wakefulness and unconsciousness.^76–78^ We investigated how dynamic stability covaries with these fluctuations in responsiveness.

We found that the degree of instability is an effective indicator of the level of responsiveness exhibited within a single session. Throughout the sessions, NHPs were presented with tones indicating that they should press a lever. To identify the periods in which the NHPs were behaviorally responsive, we computed the percent of correct lever presses within two minute windows. Responsive windows were taken to be those in which the NHP pressed the lever to more than 10% of the tones. We then computed the mean instability values from DeLASE within these windows. To illustrate this, we plotted the responsive periods (green panels) and mean instability (red curves) for two example sessions from each NHP (Figure 5A,B). The time series of instability fluctuated dramatically throughout these sessions, reflecting the periods of responsiveness exhibited by the NHPs. Notably, instability peaked during brief moments of unresponsiveness and dipped during brief moments of responsiveness (Figure 5A,B).

**Figure 5:**
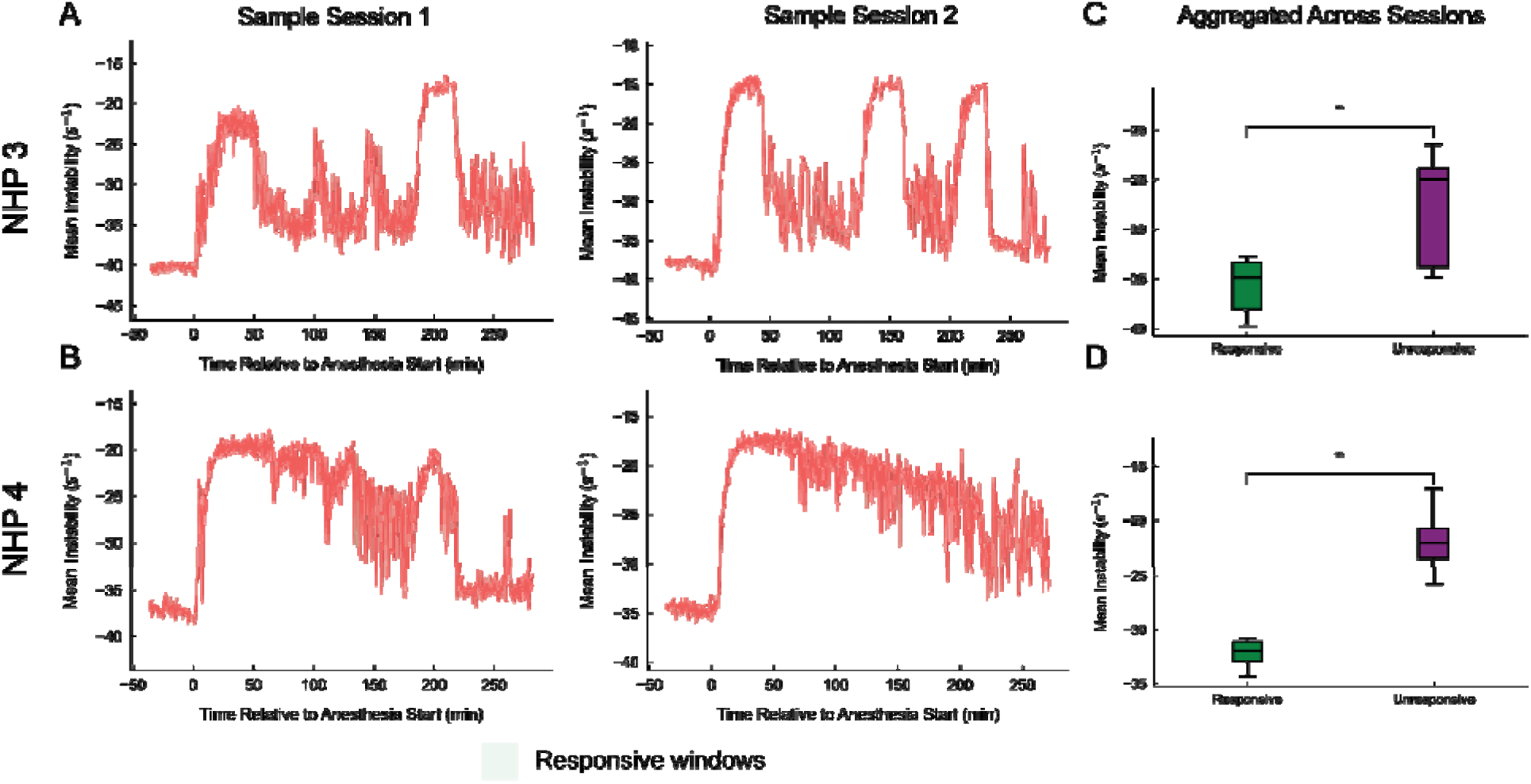
Instability tracks wakefulness within single dexmedetomidine sessions. Data are represented as mean ± SEM, with results averaged over characteristic roots for (A, B) and over sessions for each NHP for (C, D). (A, B) Two example sessions for NHP 3 (A) and NHP 4 (B). Periods in which the NHP was identified as responsive, based on the percent of successful lever presses, are plotted with green panels. Mean instability is plotted with red curves. (C, D) Mean instability within responsive and responsive periods for NHP 3 (C) and NHP 4 (D) aggregated across sessions.

To quantify this, we computed the average instability within responsive and unresponsive periods. We found that instability was greater during the unresponsive periods in both NHPs (Figure 5C,D, p < 0.01 for both NHPs, one-sided Wilcoxon signed-rank test). We note here that while we used a binary distinction for this analysis (i.e., responsive v.s. unresponsive), a more nuanced classification may yield more informative results.

## Discussion

Our results show that propofol, ketamine, and dexmedetomidine destabilize cortical dynamics. The destabilization was evident in neural responses to sensory inputs, which exhibited longer timescales during anesthesia. The effect of this would be increased low-frequency power. Indeed, cortical destabilization was correlated with increased power in the low-frequency spectrum (0.1 - 4 Hz delta band, 4 - 8 Hz theta band, 8 - 12 Hz alpha) band. All three anesthetics destabilized cortical activity to a similar degree. This suggests that, despite their different actions on different molecular receptors in different sets of neurons, there is a common signature of LOC at a more emergent level of cortical dynamics.

These results are consistent with previous work showing that, under anesthesia, cortical activity shows longer intrinsic timescales,^79–83^ a characteristic of destabilized systems.^84–87^ This matches previous results demonstrating that propofol, ketamine, and dexmedetomidine all increase low-frequency power in cortex.^6,10,12,21–24^ Prior work on cortical stability during anesthetic infusion produced contradictory results, suggesting that infusion either destabilizes^88–90^ or excessively stabilizes^91–94^ neural dynamics. Our prior work developed DeLASE, a new tool for measuring stability^30^. Cortical activity is highly nonlinear^95–99^. DeLASE takes into account nonlinear dynamics as well as our limited observations (250 or so recording sites) of a much larger system, and is computationally tractable. This revealed that propofol destabilized cortex. Our current results showing that destabilization of cortex is a common signature of LOC points to the importance of stabilization for consciousness. In fact, the relationship between stability and consciousness is not limited to anesthesia. It has been theorized that dynamic stability and consciousness are closely linked.^100–110^ When humans are presented with threshold-level sensory stimuli, neural representations are more stable when the stimuli were perceived vs not perceived.^111^ Similarly, in our dexmedetomidine sessions our NHPs fluctuated in and out of unconsciousness. Correspondingly, cortex fluctuated between stability and destabilization. This suggests that dynamic instability may serve as a sensitive marker of consciousness level. For example, it could be used, via EEG, to track level of consciousness during surgery.^112,113^

Ketamine showed unique dynamics. At first, there was a destabilization shortly after the bolus injection that induced LOC. After this initial dose began to wear off, and the NHPs began to regain consciousness, the cortex was actually more stable than during wakefulness. Correspondingly, autocorrelation showed shorter time constants, low-frequency power (XX Hz) decreased, and high-frequency power (XX Hz) increased. Although we did not study ketamine’s dose-dependent effects directly, this reinforces the observation that lower doses of ketamine have distinct effects from anesthetic doses.^16,26,114–116^ A study using a ketamine dose approximately one fourth of ours found increased stability in cortex and supports this conclusion.^117^ Also, higher doses of ketamine decrease the complexity of brain activity while lower-doses increase complexity.^94,118–126^ Complexity means richer representations, which presumably depends on a stable cortex. But not too much stability. LOC due to seizures is associated with a large increase in cortical stability (Toker ref). Low-dose ketamine has shown some promise as a treatment for mood disorders.^127–137^ Further work will be needed to determine if these treatment effects are after-effects of the cortex in a higher frequency, relatively stable state.

It has been suggested that propofol-induced destabilization is a result of the increase in inhibitory strength induced by the molecular action of propofol.^30^ An alternative hypothesis, not inconsistent with our results, is that the destabilization is related to the gating of bottom-up sensory inputs to cortex. Strong inputs can stabilize otherwise unstable dynamics.^138–141^ Anesthesia may disrupt the relay of sensory input from subcortical to cortical regions.^21,142–148^ In particular, it has been shown to disrupt the progression of these inputs to higher level cortices.^149–151^ This is consistent with theories of consciousness that propose formation of meta ensembles that coordinate cortex on a large scale.^104,152–159^ Thus the observed destabilization may reflect a lack of input-induced stabilization as a result of disrupted sensory inputs. This hypothesis is supported by the fact that deep-brain stimulation can induce arousal from anesthetic-induced unconsciousness.^12,160–163^

Leveraging high-resolution intracortical recordings and dynamical systems analysis, we provide evidence for destabilization as a convergent effect of different anesthetics. This puts forward destabilization as a potentially common signature of loss of consciousness among anesthetics. This points to an optimal range of dynamic stability as a prerequisite for consciousness.

## Acknowledgements

The authors would like to acknowledge Leo Kozachkov for thoughtful and helpful discussions. E.K.M. acknowledges funding from ONR MURI N00014-23-1-2768, NIMH 1R01MH131715-01, NIMH R01MH11559, The Simons Center for the Social Brain, The Freedom Together Foundation, and The Picower Institute for Learning and Memory. I.R.F. acknowledges funding from The National Science Foundation Computer and Information Science and Engineering Directorate, The Simons Collaboration on the Global Brain, and The McGovern Institute at MIT. E.N.B. acknowledges funding from The Freedom Together Foundation, and The Picower Institute for Learning and Memory. Y.I. acknowledges funding from NIH 1R21AG077275-01A1.

## Supplementary Figures

**Figure S1:**
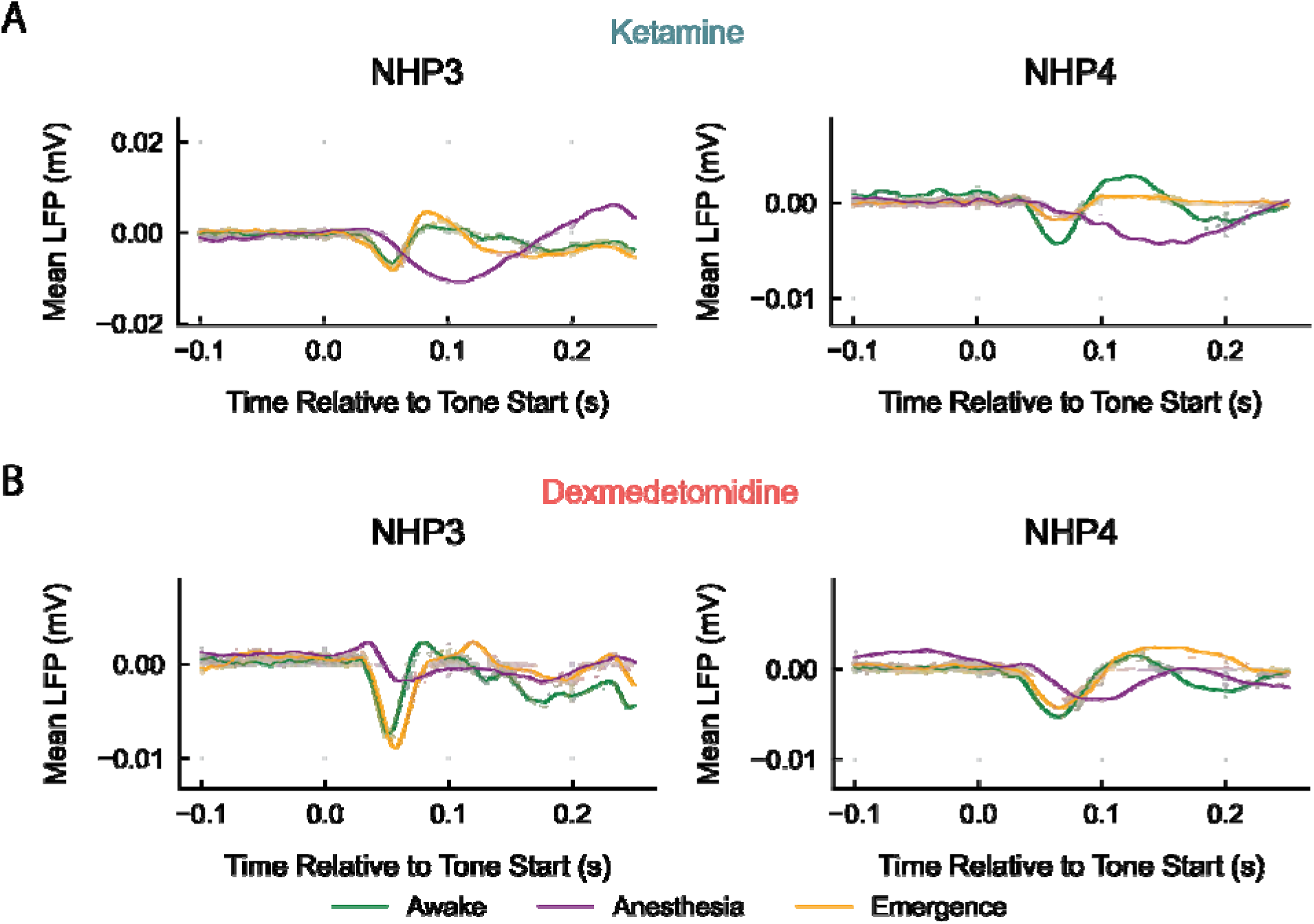
Temporally zoomed sensory responses from ketamine and dexmedetomidine. Data are represented as mean ± SEM, with results averaged over sessions for each NHP. (A, B) The zoomed in LFP response to auditory stimuli from Figure 3 for ketamine (A) and dexmedetomidine (B). Response was averaged across electrodes and trials within a session, then the mean was computed across sessions. Translucent curves correspond to response averages from single sessions. Green curve is the Awake epoch, purple is the Anesthesia epoch, and orange is the Emergence epoch. Auditory oddball stimuli consisted of five 150 ms spaced by 50ms, of which the first 250 ms is depicted here.

**Figure S2:**
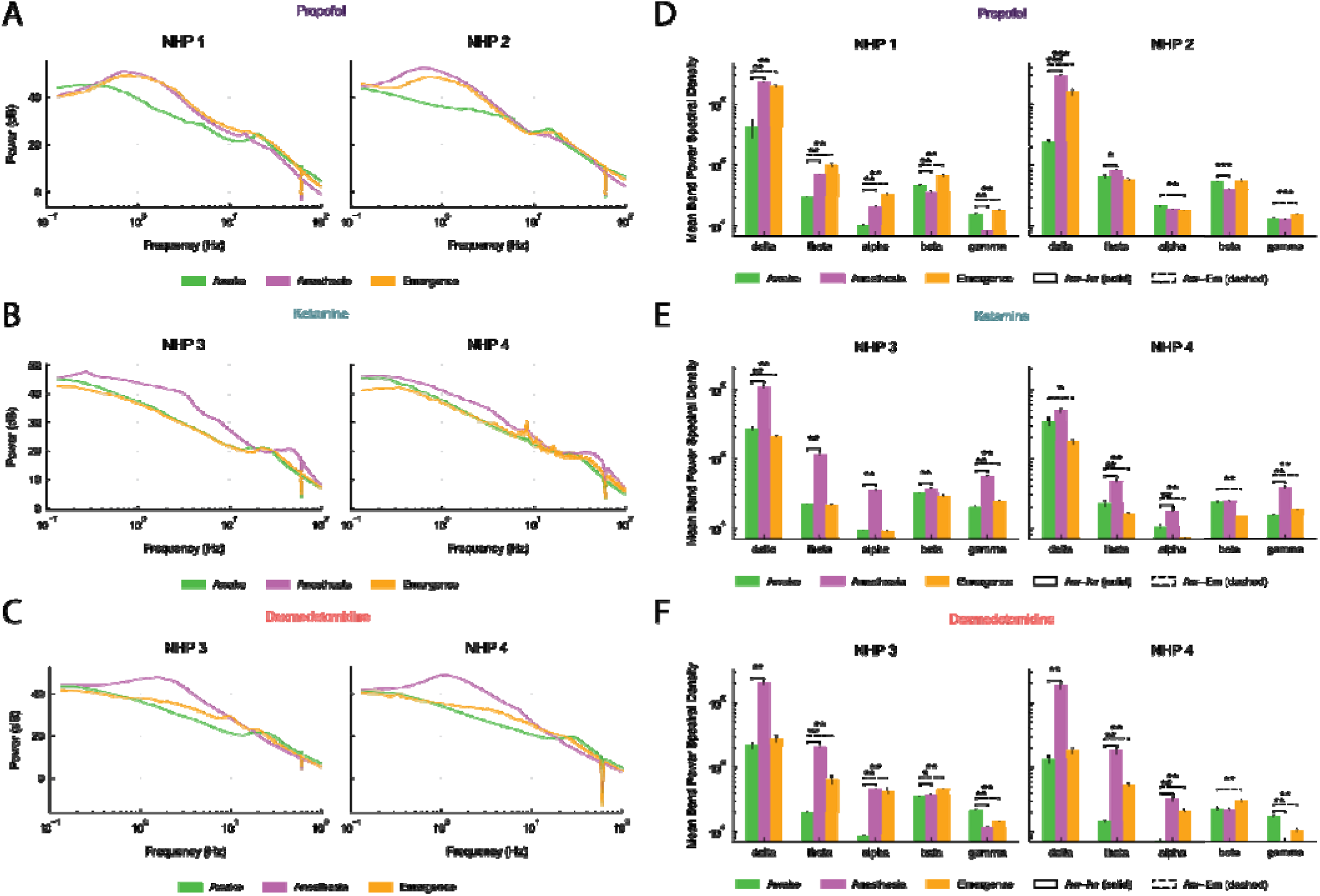
Power spectral analysis across anesthetic agents and epochs. Data are represented as mean ± SEM, with results averaged over sessions for each NHP. (A, B, C) The mean power spectral densities, averaged across sessions and within epochs, for propofol (A), ketamine (B), and dexmedetomidine (C). (D, E, F) Comparison of band power between epochs for propofol (A), ketamine (B) and dexmedetomidine (C). Statistics are computed over sessions.

**Figure S3:**
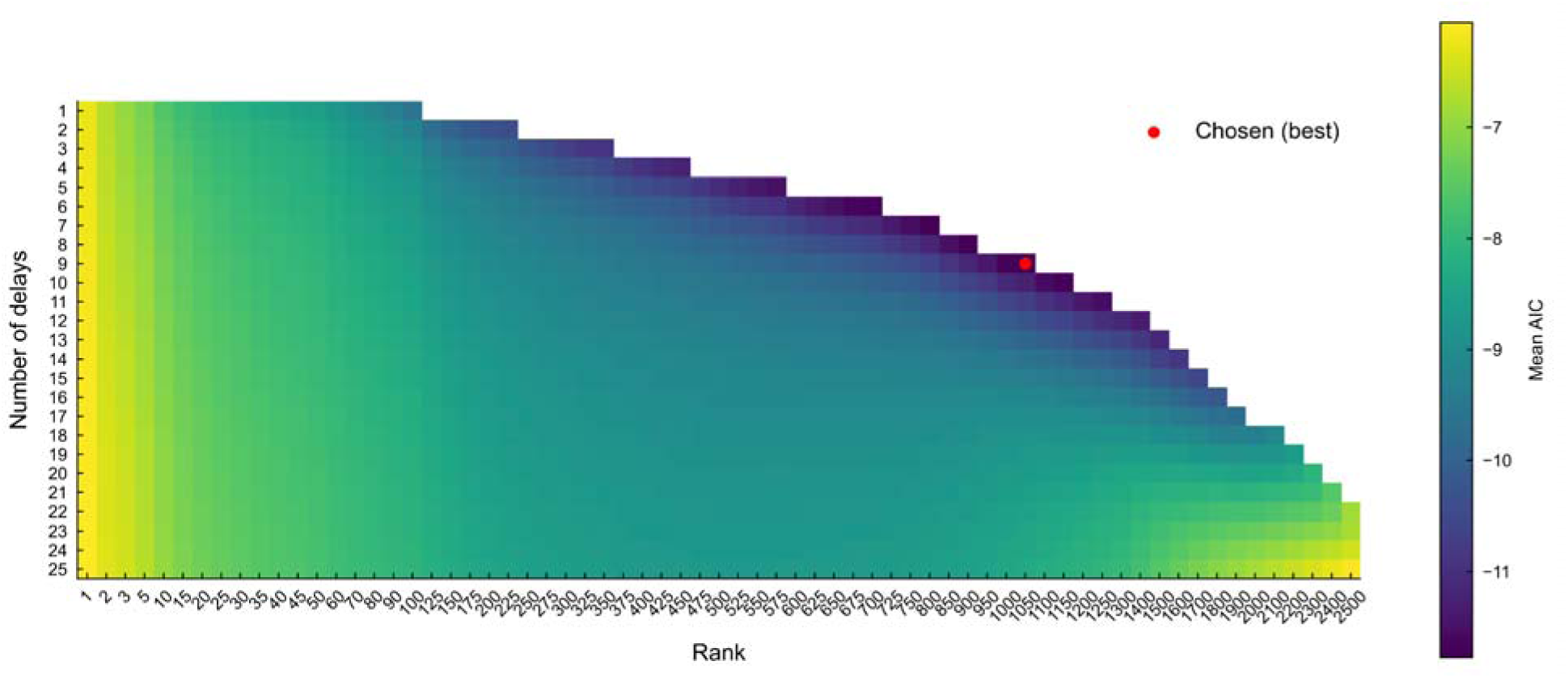
Sample grid search results for a single dexmedetomidine session. The mean AIC on test-set one-step prediction on 32 15 second windows a single session, chosen such that there were 4 windows per subsection of the session (see methods). The y-axis is the minimum number of delay embedding coordinates, effectively determining the number of lags. The x-axis is the rank of the dynamics, or number of independent temporal modes, extracted through SVD on the delay embedding matrix that are used for fitting the dynamical model. Color indicates Akaike Information Criterion (AIC) normalized to the number of predicted values. AIC quantifies the balance between high quality model prediction (as measured by one-step prediction error on test data) with the number of model parameters. Models are penalized for having larger numbers of parameters, such that the optimal model of minimal complexity can be obtained.

